## Supplemental methods for "Ligand Discrimination in Immune Cells: Signal Processing Insights into Immune Dysfunction in ER+ Breast Cancer"

### Supplementary material

### Table S1. Breast cancer patient characteristics.

| SAMPLE ID | AGE AT DX | ER | PR | HER2 | KI67 (%) | TMR GRADE | T | N | M | STAGE | SEX |
|---|---|---|---|---|---|---|---|---|---|---|---|
| **19186-2** | 41 | + | + | - | 25 | 2 | 1b | 0 | X | Ia | F |
| **19186-3** | 42 | + | + | - | 10 | 1 | 1b | 0 | X | Ia | F |
| **19186-4** | 48 | + | + | - | 1 | 2 | 1b | 0 | X | Ia | F |
| **19186-8** | 71 | + | + | - | 5-10 | 1 | 1B | 1a | X | IIa | F |
| **19186-10** | 48 | + | + | - | 10 | 2 | 2 | 0 | X | IIa | F |
| **19186-14** | 51 | + | + | - | 25 | 2 | 2 | 1a | X | IIb | F |
| **19186-15** | 73 | + | + | - | 5 | 0 | 3 | 0 | X | IIb | F |
| **21368-003** | 35 | + | + | - | 20-30 | 2 | 2 | 1 | X | IIb | F |
| **21368-004** | 65 | + | + | - | 10-15 | 2 | mT2 | 0 | X | IIa | F |
| **21368-005** | 49 | + | + | - | 5-10 | 2 | 1c | 0 | X | Ia | F |
| **21368-007** | 57 | + | + | - | <3 | 2 | 1b | 0 | X | Ia | F |
| **21368-008** | 76 | + | + | - | 3-5 | 1 | 1c | 0 | X | Ia | F |
| **21368-010** | 40 | + | + | - | 10 | 1 | 1b | 0 | X | Ia | F |
| **21368-011** | 73 | + | + | - | 20 | 2 | 2 | 0(i+) | X | IIa | F |
| **21368-012** | 47 | + | + | - | 30 | 2 | 1c | 1 | X | IIa | F |
| **21368-014** | 74 | + | + | - | Unknown | 1 | 1a | 0 | X | Ia | F |
| **21368-015** | 59 | + | + | - | 6 | 2 | 1c | 0 | X | Ia | F |
| **21368-018** | 75 | + | + | - | Unknown | 2 | 1b | 1a | X | IIa | F |

### Table S2. Healthy donor characteristics.

| SAMPLE ID | AGE | SEX |
|---|---|---|
| **HD1** | 66 | F |
| **HD2** | 79 | F |
| **HD3** | 54 | F |
| **HD4** | 70 | F |
| **HD5** | 35 | F |
| **HD6** | 43 | F |
| **HD7** | 44 | F |
| **HD8** | 45 | F |
| **HD9** | 52 | F |
| **HD10** | 56 | F |
| **HD11** | 63 | F |
| **HD12** | 58 | F |
| **HD13** | 26 | F |
| **HD14** | 28 | F |
| **HD15** | 30 | F |
| **HD16** | 34 | F |
| **HD17** | 52 | F |
| **HD18** | 60 | F |
| **HD19** | 61 | F |
| **HD20** | 62 | F |
| **HD21** | 68 | F |
| **HD22** | 69 | F |
| **HD23** | 70 | F |
| **HD24** | 41 | F |
| **HD25** | 26 | F |
| **HD26** | 52 | F |
| **HD27** | 59 | F |
| **HD28** | 62 | F |
| **HD29** | 69 | F |
| **HD30** | 70 | F |
| **HD31** | 67 | F |
| **HD32** | 47 | F |

**Flow cytometry gating strategy**

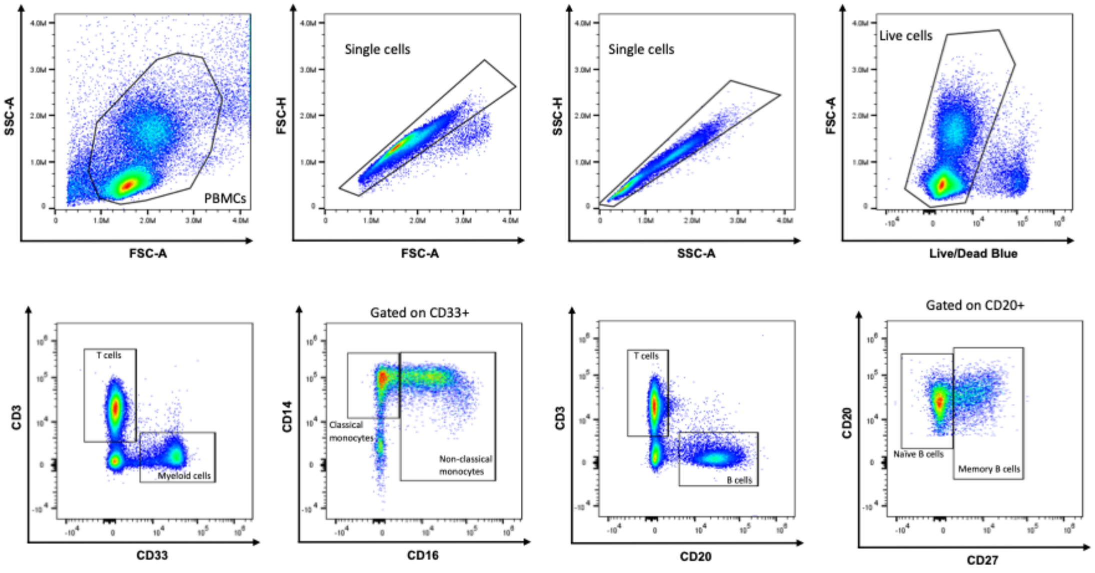

**Figure S1.** Identification of PBMCs, single cells, live cells, and T, myeloid, monocyte, and B cell populations.

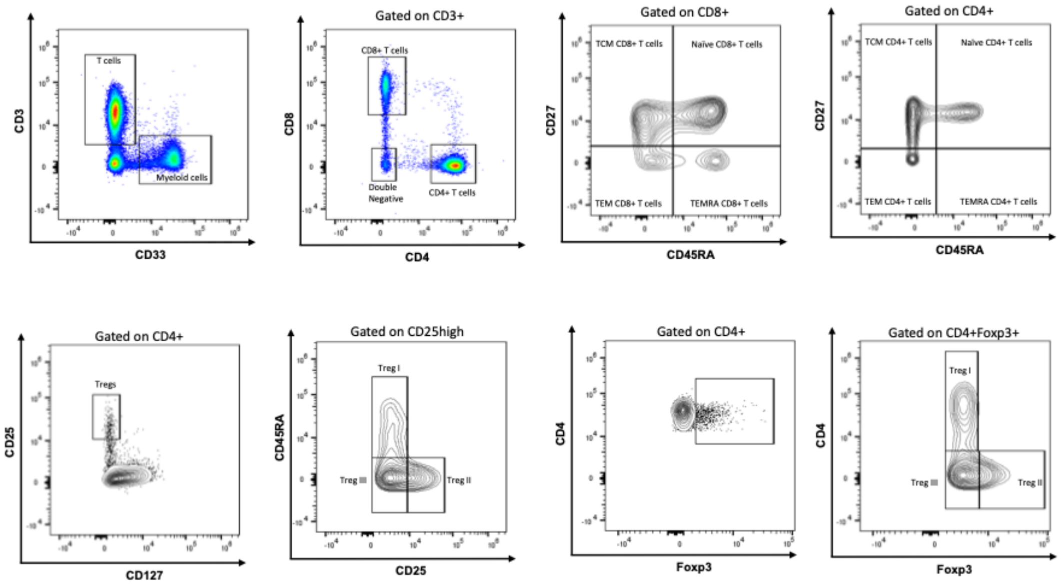

**Figure S2.** Identification of T cell subsets.

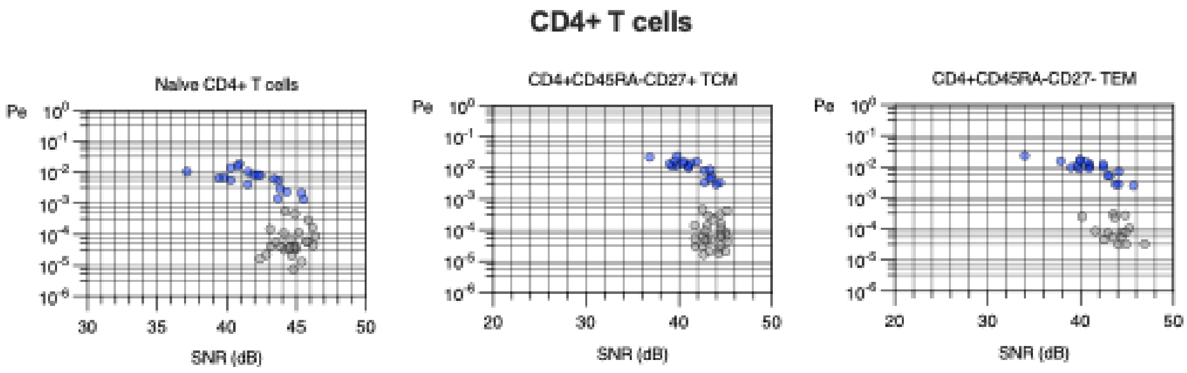

**Figure S3.** Signal error and SNR for CD4+ T cell subsets.

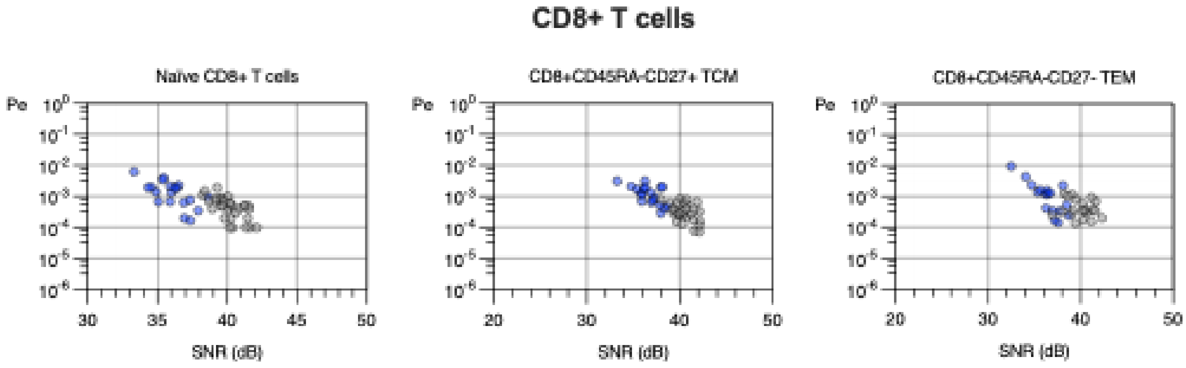

**Figure S4.** Signal error and SNR for CD8+ T cell subsets.

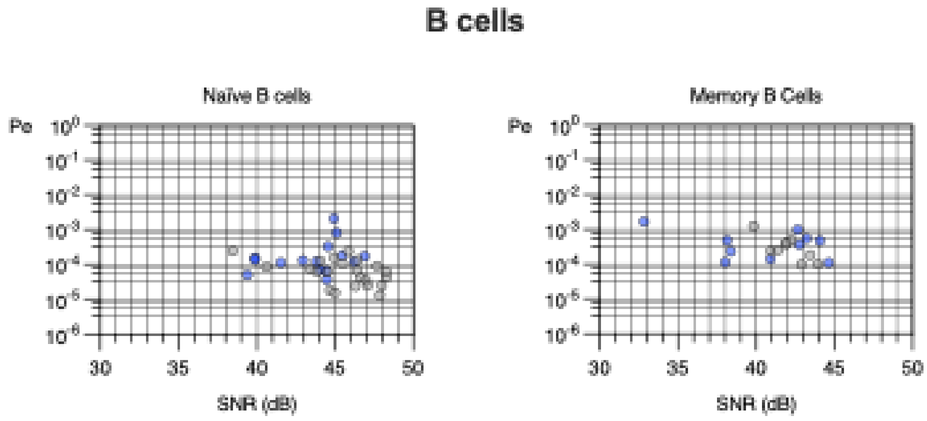

**Figure S5.** Signal error and SNR for B cell subsets.

### Confusion matrix analysis

As shown in the main text, we examined pair-wise error rates by cell type for all breast cancer samples (N=19), represented as a confusion matrix. Darker shades of grey correspond to higher error rates, or equivalently lower values of $-\ln(P_e)$, shown for each pair of true-detected signals. A value of “Inf” indicates the probability of error is zero, and therefore the quantity $-\ln(P_e)$ is undefined and set to infinity. Values along the diagonal are not relevant to this analysis. Confusion matrix analysis reveals IL-2 and IL-4 as being a pair of cytokines that may partially explain the high error rates observed in CD4+ T cells as compared to other immune cell subtypes.

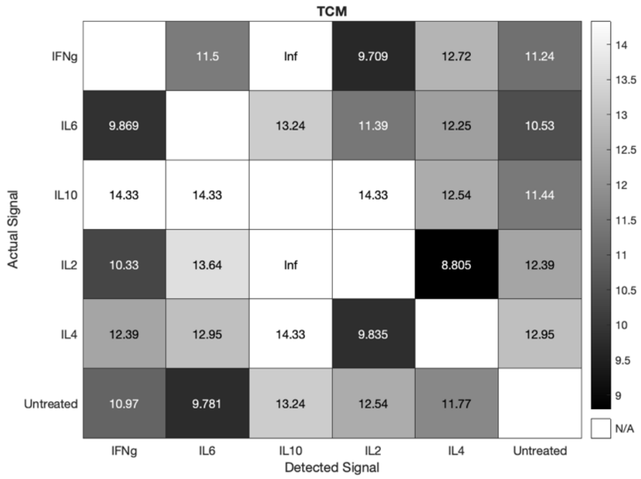

**Figure S6.** Confusion matrix for CD4+ TCM cells given by $-\ln(P_e)$. The most likely signal confusion is between IL-4 and IL-2 cytokine stimulation, followed by INF-γ and IL-2.

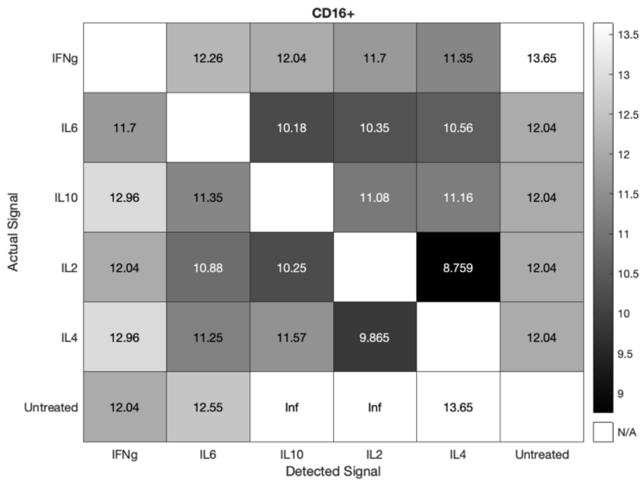

**Figure S7.** Confusion matrix for CD16+ NK cells given by $-\ln(P_e)$. The most likely signal confusion is between IL-4 and IL-2 cytokine stimulation.

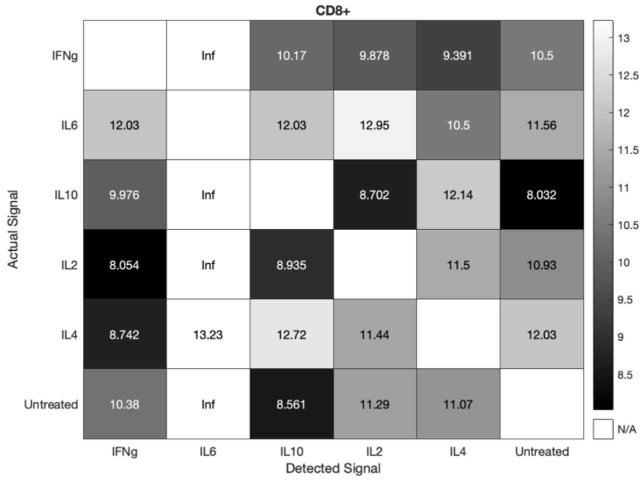

**Figure S8.** Confusion matrix for CD8+ T cells given by $-\ln(P_e)$. The most likely signal confusion is between IL-10 and IL-2 cytokine stimulation, followed by INF-γ and IL-4.

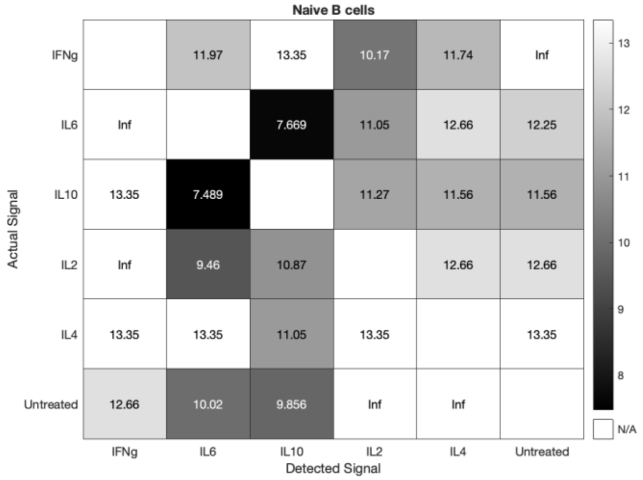

**Figure S9.** Confusion matrix for naïve B cells given by $-\ln(P_e)$. The most likely signal confusion is between IL-6 and IL-10 cytokine stimulation, followed by IL-6 and IL-2.

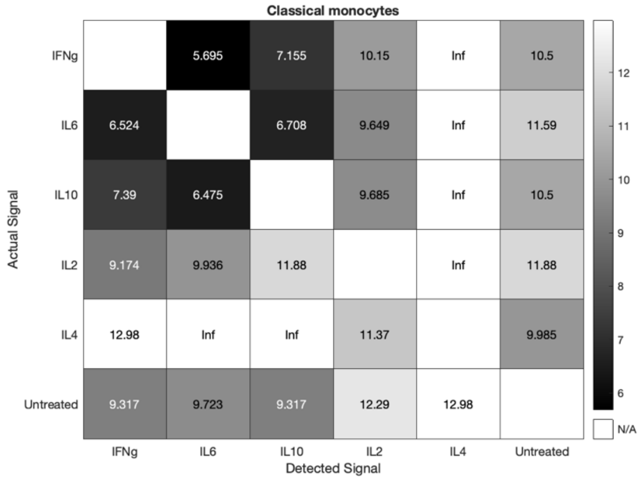

**Figure S10.** Confusion matrix for classical monocytes given by $-\ln(P_e)$. The most likely signal confusion is between IL-6 and INF-γ cytokine stimulation, followed by IL-6 and IL-10.

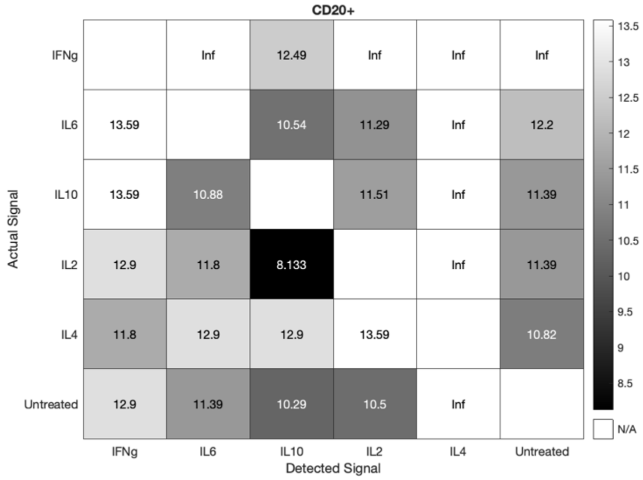

**Figure S11.** Confusion matrix for CD20+ B cells given by $-\ln(P_e)$. The most likely signal confusion is between IL-10 and IL-2 cytokine stimulation.

**Receptor expression correlation analysis**

To interrogate the potential relationship between receptor expression and computed probability of error or signal-to-noise ratio, we performed correlation analysis on a set of relevant cell surface receptors: CD132 (IL2Rg), CD122 (IL2Rb), CD124 (IL4Ra), CD130 (IL6Rb), CD126 (IL6Ra), CD210 (IL10R), CD119 (INFgR1), PDL1, and PD1. We analyzed both the MFI of the expression level and a receptor SNR (rSNR), computed from the log transformed MFI data (see equation 1 in the main text). Results for MFI and rSNR were similar. We show rSNR results in the tables and figures below. No single receptor or combination of receptors were sufficient to explain Pe, SNR, or observed differences in Pe or SNR between healthy donors and ER+ breast cancer samples.

| | Probability of error (Pe) | | | | | | | | | |
|---|---|---|---|---|---|---|---|---|---|---|
| Receptor | | | | | | | | | | |
| HD | CD4+ Naïve | CD4+TCM | CD4+TEM | CD8 + Naïve | CD8+TCM | CD8+TEM | Classical Monocytes | NK Cells | Naïve B cells | Memory B cells |
| CD132 (IL2Rg) | | | | | ** | | | | | |
| CD122 (IL2Rb) | | | | | | | | | | |
| CD124 (IL4Ra) | | | | | | | | ** | | |
| CD130 (IL6Rb) | | | | | * | | * | | | |
| CD126 (IL6Ra) | | | | * | | | | | | |
| CD210 (IL10R) | | | | | | | | | | |
| CD119 (IFNgR1) | | **** | | | | | | *** | | |
| PDL1 | | | | | | | | | | |
| PD1 | | | | | * | | | *** | | |
| BC | CD4+ Naïve | CD4+TCM | CD4+TEM | CD8 + Naïve | CD8+TCM | CD8+TEM | Classical Monocytes | NK Cells | Naïve B cells | Memory B cells |
| CD132 (IL2Rg) | | | | | | | | * | | |
| CD122 (IL2Rb) | ** | | | | | | | | | |
| CD124 (IL4Ra) | * | | | | * | | | | | |
| CD130 (IL6Rb) | | | | | | | | | | |
| CD126 (IL6Ra) | * | | | | * | | | | | |
| CD210 (IL10R) | | | | | | | | | | |
| CD119 (IFNgR1) | | | | | | | | | | |
| PDL1 | | | | | | * | * | | | |
| PD1 | | | | | | | * | | | |
| | | | positive correlation | | | | | | | |
| | | | negative correlation | | | | | | | |

**Figure S12. Probability of error receptor expression correlation analysis.** Receptor expression was correlated with Pe for each cell type in all HD samples (n=32, top) and BC samples (n=19, bottom). Statistically significant positive correlations are shown in green and negative correlations are shown in red ($* p < 0.05$, $** p < 0.01$, $*** p < 0.001$, $**** p < 0.0001$).

| | Signal-to-noise ratio (SNR) | | | | | | | | | |
|---|---|---|---|---|---|---|---|---|---|---|
| Receptor | | | | | | | | | | |
| HD | CD4+ Naïve | CD4+ TCM | CD4+ TEM | CD8 + Naïve | CD8+ TCM | CD8+ TEM | Classical Monocytes | NK Cells | Naïve B cells | Memory B cells |
| CD132 (IL2Rg) | | | | | * (green) | | | * (green) | | * (red) |
| CD122 (IL2Rb) | | | | | | | | * (red) | | |
| CD124 (IL4Ra) | | | | | * (red) | | | | | |
| CD130 (IL6Rb) | | | | | | | | | | * (green) |
| CD126 (IL6Ra) | | | | | | | | | | (green) |
| CD210 (IL10R) | | | | | | | | | | |
| CD119 (IFNgR1) | | | * (green) | | | | | | | |
| PDL1 | | | | | | | | | | |
| PD1 | | | | | | | | | | |
| BC | CD4+ Naïve | CD4+ TCM | CD4+ TEM | CD8 + Naïve | CD8+ TCM | CD8+ TEM | Classical Monocytes | NK Cells | Naïve B cells | Memory B cells |
| CD132 (IL2Rg) | | | * (red) | * (red) | | | *** (red) | | | * (green) |
| CD122 (IL2Rb) | | | | | | | ** (green) | ** (green) | | |
| CD124 (IL4Ra) | | | | | * (green) | | | *** (green) | | |
| CD130 (IL6Rb) | | | | | ** (red) | | ** (green) | | | *** (green) |
| CD126 (IL6Ra) | | | | | | | | | | |
| CD210 (IL10R) | | | * (green) | * (red) | | | ** (red) | | | |
| CD119 (IFNgR1) | | | | * (green) | ** (green) | | ** (green) | ** (green) | * (green) | *** (green) |
| PDL1 | | | | | | | | * (red) | | |
| PD1 | | | | | | | | | | |
| | | (green) | positive correlation | | | | | | | |
| | | (red) | negative correlation | | | | | | | |

**Figure S13. Signal-to-noise receptor expression correlation analysis.** Receptor expression was correlated with SNR for each cell type in all HD samples (n=32, top) and BC samples (n=19, bottom). Statistically significant positive correlations are shown in green and negative correlations are shown in red ($* \; p < 0.05$, $** \; p < 0.01$, $*** \; p < 0.001$, $**** \; p < 0.0001$).

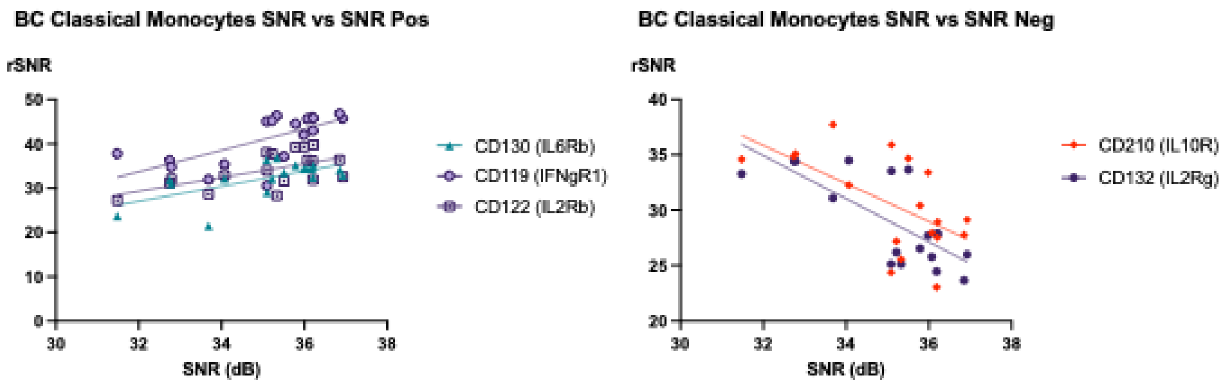

**Figure S14. Correlation analysis of receptor expression rSNR on classical monocytes with overall SNR in breast cancer samples.** Statistically significant correlations were found in expression levels of CD130, CD119, CD122, CD210, and CD132 on classical monocytes in breast cancer samples (n=19). CD130, CD119, and CD122 were positively correlated, and CD210 and CD132 were negatively correlated. Because overall SNR is computed by integrating all pSTAT responses from all cytokine stimulations, these correlations do not explain differences observed between healthy donors and BC samples; moreover, classical monocytes show the lowest SNR in HD and BC samples for any immune cell subtype.

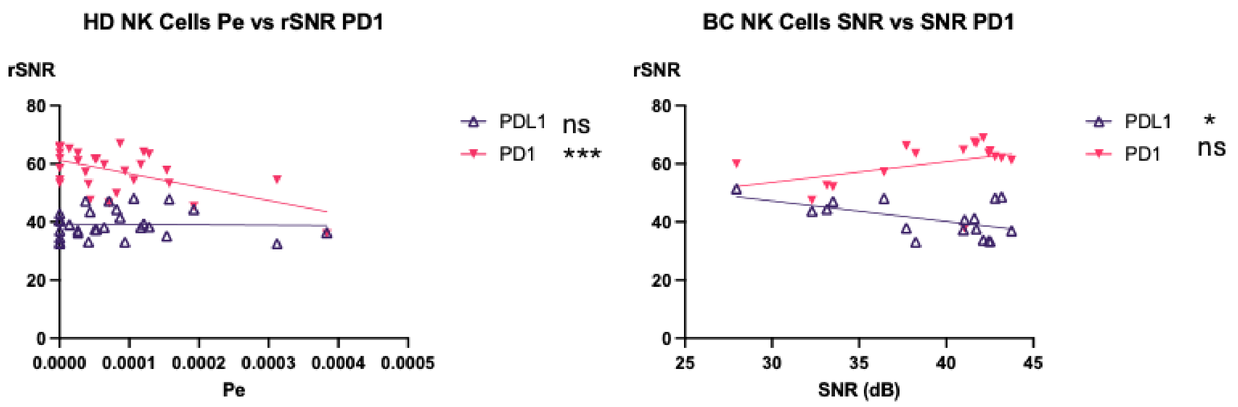

**Figure S15. Correlation analysis of PDL and PD1 expression rSNR on NK cells with Pe and overall SNR in healthy donors and breast cancer samples.** Because of the clinical relevance of PDL1 and PD1 expression on immune cells, we examined the correlation of PD(L)1 expression levels in NK cells for healthy donors (n=32) and breast cancer (n=19) samples. Neither PDL1 nor PD1 were consistently correlated with Pe or SNR in either healthy donors or breast cancer samples. The significance of the correlation is driven by large outliers in Pe and small outliers in SNR.
